# G2019S selective LRRK2 kinase inhibitor abrogates mitochondrial DNA damage

**DOI:** 10.1101/2022.11.30.517979

**Authors:** Nicholas Pena, Claudia P. Gonzalez-Hunt, Rui Qi, Carrolee Barlow, Natalie F. Shanks, Holly J. Carlisle, Laurie H. Sanders

## Abstract

Pathogenic mutations in *LRRK2* cause Parkinson’s disease (PD). The G2019S variant is the most common, which results in abnormally high kinase activity. Compounds that target LRRK2 kinase activity are currently being developed and tested in clinical trials. We recently found that G2019S LRRK2 causes mitochondrial DNA (mtDNA) damage and treatment with multiple classes of LRRK2 kinase inhibitors at concentrations associated with dephosphorylation of LRRK2 reversed mtDNA damage to healthy control levels. Because maintaining the normal function of LRRK2 in heterozygous G2019S *LRRK2* carriers while specifically targeting the G2019S LRRK2 activity could have an advantageous safety profile, we explored the efficacy of a G2019S mutant selective LRRK2 inhibitor to reverse mtDNA damage in G2019S LRRK2 models and patient cells relative to non-selective LRRK2 inhibitors. Potency of LRRK2 kinase inhibition by EB-42168, a G2019S mutant LRRK2 kinase inhibitor, and MLi-2, a nonselective inhibitor, was determined by measuring phosphorylation of LRRK2 at Ser935 and/or Ser1292 using quantitative western immunoblot analysis. The Mito DNA_DX_ assay, a novel system that allows for the accurate real-time quantification of mtDNA damage in a 96-well platform, was performed in parallel. We confirmed that EB-42168 selectively inhibits LRRK2 phosphorylation on G2019S LRRK2 relative to wild-type LRRK2. On the other hand, MLi-2 was equipotent for wild-type and G2019S LRRK2. Acute treatment with EB-42168 inhibited LRRK2 phosphorylation and also restored mtDNA damage to healthy control levels. Precision medicine is a common approach in modern day cancer research that is not yet routinely applied to neurodegenerative diseases. Abrogation of mtDNA damage with mutant selective tool inhibitor EB-42168 demonstrates the promise of a precision medicine approach for LRRK2 G2019S PD. Levels of mtDNA damage may serve as a potential pharmacodynamic biomarker of altered kinase activity that could be useful for small molecule development and clinical trials.

## INTRODUCTION

Parkinson’s disease (PD) is the most common age-related movement neurodegenerative disorder. This chronic disease is characterized by progressive motor disability, non-motor symptoms and decreased quality of life. Therapeutic strategies currently available rely on dopamine replacement, but do not slow or stop the progression of the disease and can lead to motor complications. While these drugs may be useful in managing symptoms, there are no disease-modifying therapies that target the underlying pathogenic mechanisms of disease; thereby this remains a significant and urgent unmet medical need for PD patients. In 2004, coding variants were first identified in *Leucine-rich repeat kinase 2* (LRRK2), with missense mutations in *LRRK2* now established as the most common genetic cause of autosomal-dominant PD [1-5]. LRRK2 is a large and complex protein, with two functional enzymatic domains – a Ras-like GTPase and a serine-threonine kinase domain. The most frequent pathogenic *LRRK2* mutation is the Gly2019Ser (G2019S) *LRRK2* variant, which results in a modest increase in kinase activity [6-8]. Interestingly, in addition to the *LRRK2* G2019S variant, all pathogenic missense *LRRK2* mutations also augment kinase activity, displaying higher levels of the autophosphorylation Ser1292 residue of LRRK2 [9-12]. Therefore, the toxic gain-of-function of LRRK2 kinase activity is strongly implicated as the cause of pathogenicity [13]. Consistent with these findings, a neuroprotective effect of LRRK2 kinase inhibitors has been demonstrated in PD-relevant cell and rodent models [14]. Thus, LRRK2 represents a promising therapeutic target for disease modification and several LRRK2 kinase inhibitors are in clinical development and/or trials.

Recently, the results of testing DNL201, a CNS-penetrant, selective, ATP-competitive, small-molecule LRRK2 kinase inhibitor in early phase human clinical trials were reported [15]. Importantly, DNL201, in humans was able to cross the blood brain barrier, based on measurements of inhibitor concentrations in cerebrospinal fluid. Concomitant blood-based markers demonstrated a dose-dependent inhibition of LRRK2 kinase by DNL201 [15]. However, on-target safety liabilities for LRRK2 inhibitors have been identified in preclinical models. Several studies have shown lung and/or kidney dyshomeostasis in rodent and nonhuman primate models either lacking LRRK2 or following treatment with LRRK2 kinase inhibitors, some of which may be reversible [16-26]. Of note, lung function did not seem to be impacted at the DNL201 doses tested in single-ascending dose or multiple-ascending dose (10 days) cohorts in healthy volunteers or in patients with PD. Longer-term monitoring will be critical to assess safety of chronic dosing with LRRK2 kinase inhibitors on lung and kidney function, as well as in PD patients with clinically significant history of pulmonary and kidney disease, which are often co-morbidities in this population. Heterozygous loss-of-function variants at the *LRRK2* locus do not increase PD risk and have no apparent overt deleterious health consequences [27, 28]. However, somatic *LRRK2* mutations in breast cancer are associated with high-risk features and reduced patient survival [29], consistent with a potential risk for lung adenocarcinoma with reduced LRRK2 levels [30], emphasizing the concern of targeting LRRK2 in humans. Additionally, since the majority of PD patients carrying the LRRK2 G2019S variant are heterozygous, a precision medicine approach of selectively reducing pathogenic LRRK2 kinase activity while sparing LRRK2 physiological function could be efficacious while offering safety advantages to patients with PD.

Measuring endogenous LRRK2 kinase activity by monitoring LRRK2 autophosphorylation at serine 1292 (pSer1292) has been particularly challenging, but can be robustly detected in overexpressed cellular models or with a fractionation-based enrichment technique in G2019S LRRK2 but not wild-type tissue [11, 31, 32]. The phosphorylation of downstream substrates, including Rab GTPase substrates, are difficult to measure reliably due to low stoichiometry [4, 5, 33, 34]. The disease relevance of phospho-substrates of LRRK2, such as Rab10, is unknown, and paradoxically, Rab10 phosphorylation is not elevated with the PD LRRK2 G2019S variant [35-38]. The indirect but kinase conformation dependent phosphorylation site at serine 935 (pSer935) is most widely used for measuring LRRK2 kinase inhibition, despite not faithfully reflecting LRRK2 protein kinase activity [12, 39, 40]. Further optimizing and developing tools to measure endogenous LRRK2 kinase activity and inhibition *in vivo*, is a critical unmet need for future clinical trials. Mitochondrial function is significantly impacted in PD [41]. Importantly, mitochondrial DNA (mtDNA) homeostasis is disrupted in both idiopathic and familial PD cases, including those with *LRRK2* mutations, and are associated with mtDNA damage [42-49]. We found that increased mtDNA damage in PD patient-derived immune cells harboring the G2019S LRRK2 mutation was abrogated following treatment with multiple classes of LRRK2 kinase inhibitors and correlated with measures of LRRK2 dephosphorylation [43, 48]. Therefore, measurement of mtDNA damage may serve as a surrogate for LRRK2 kinase activity and consequently of kinase inhibitor activity. Recently LRRK2 kinase inhibitors with significant selectivity for mutant (G2019S) LRRK2 compared to wild-type LRRK2 have been discovered, but it is unknown if these compounds have similar effects on G2019S LRRK2 dependent mtDNA damage [20, 50-52]. Thus, we have now extended these studies and determined whether mtDNA damage levels might serve as a biomarker for selective G2019S-LRRK2 kinase inhibition.

The aim of the present study was to compare the highly selective G2019S LRRK2 inhibitor (EB-42168) with the non-selective LRRK2 inhibitor (MLi-2) on mtDNA damage levels in LRRK2 models and patient cells. LRRK2 phosphorylation was examined in parallel to establish target engagement and compare these biomarker outcomes in response to the selective G2019S LRRK2 inhibitor, EB-42168. Measuring mtDNA damage together with LRRK2 phosphorylation is an innovative biomarker approach to determine LRRK2 kinase activity and may be helpful in considering the drug efficacy of compounds targeting hyperactive kinase activity in G2019S LRRK2 PD patients in the context of a clinical trial.

## RESULTS

### PD-linked *LRRK2* G2019S variant leads to increased kinase activity and mtDNA damage

LRRK2 kinase activity was assessed by measuring the relative levels of autophosphorylation at pSer1292 by immunoblot in HEK293 cells stably transfected with human LRRK2 or the G2019S variant of human LRRK2 (named WT-LRRK2 and G2019S-LRRK2, respectively) [10]. pSer1292 was increased ∼6 fold in G2019S-LRRK2 cells compared to WT-LRRK2, consistent with previous reports (Fig 1A,B, [20]). Levels of both pSer935 and total LRRK2 were similar between the WT-LRRK2 and G2019S-LRRK2 expressing cell lines (Fig 1C-E). The similar expression levels of LRRK2 between the two cell lines allowed for a direct comparison of the two genotypes on mtDNA damage phenotypes. G2019S-LRRK2 induced a higher level of mtDNA damage relative to WT-LRRK2 expressing cells (Fig. 1F), without any changes in the steady state of mtDNA copy number (Fig. 1G). These results are consistent with our prior study showing that mtDNA damage was increased in primary midbrain neurons overexpressing G2019S LRRK2 compared to wild-type LRRK2, a kinase dead LRRK2 mutant or the GFP expressing control [43].

**Fig 1.**
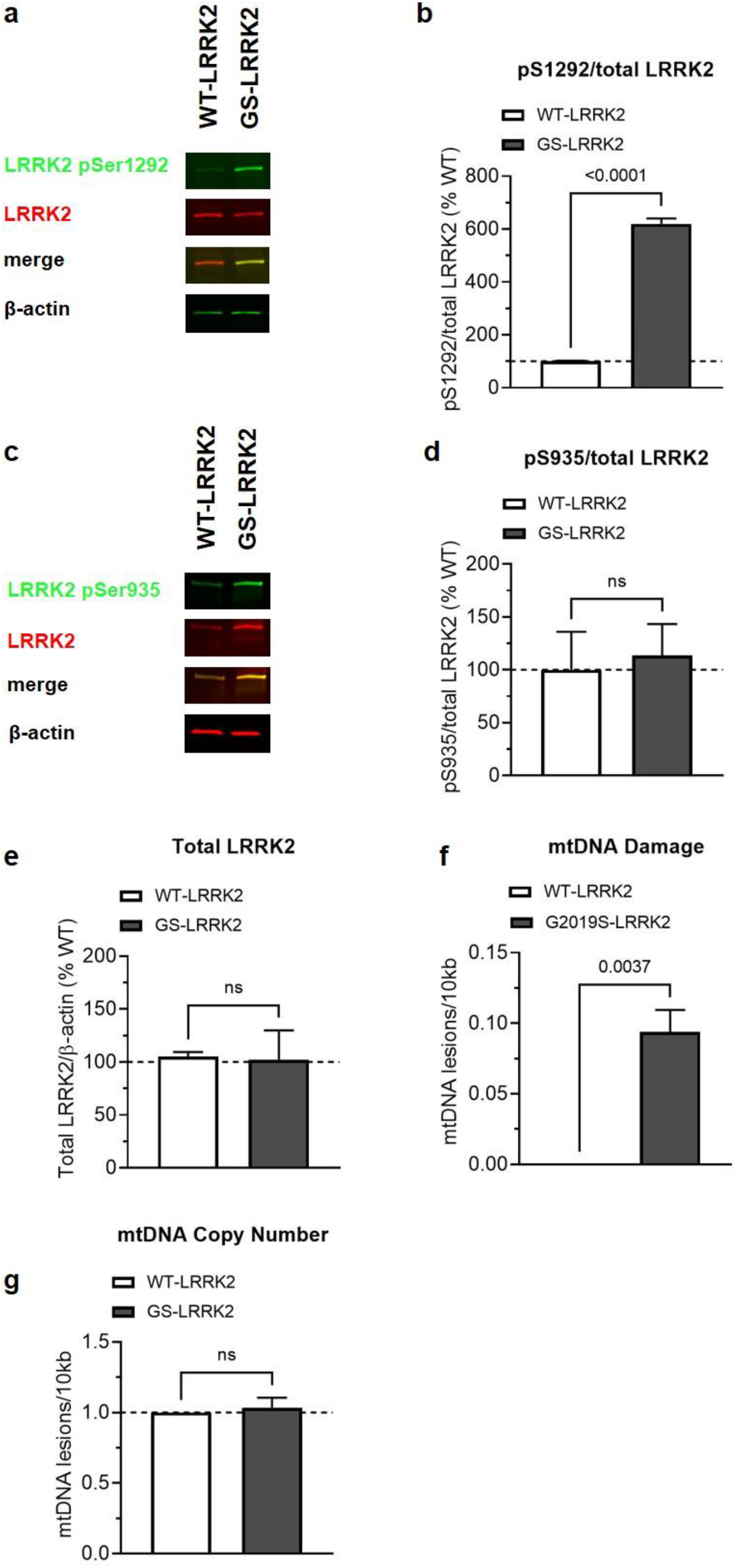
Analysis of LRRK2 and mtDNA damage in HEK293 cells lines stably overexpressing either WT-LRRK2 or G2019S-LRRK2. **a** Representative western blot of WT-LRRK2 or G2019S-LRRK2 overexpressing cells show expression of LRRK2 pSer1292 and full-length LRRK2. β-actin was blotted as a loading control. **b** Quantification of western blots demonstrate ∼ 6-fold increase of LRRK2 pSer1292 in G2019S-LRRK2 compared to WT-LRRK2 expressing cells. **c** Representative western blots of WT-LRRK2 or G2019S-LRRK2 overexpressing cells show expression of LRRK2 pSer935 and full-length LRRK2. β-actin was blotted as a loading control. **d** Quantification of western blots demonstrate no difference of LRRK2 pSer935 levels between G2019S-LRRK2 and WT-LRRK2 expressing cells. **e** Quantification of western blots demonstrate no difference of LRRK2 protein levels between the two cell lines. **f** Mitochondrial DNA damage was increased in G2019S-LRRK2 relative to WT-LRRK2 expressing cells. **g** The differences in mtDNA damage between the cell lines were not attributable to changes in steady state mtDNA levels. Data are presented as mean ± SEM. (*P<0.01 determined by unpaired t test and paired t test for total LRRK2 levels). G2019S-LRRK2 = GS-LRRK2, wild-type LRRK2 = WT-LRRK2, non-significant = ns

### EB-42168 demonstrates selective inhibition of LRRK2 phosphorylation in G2019S-LRRK2 relative to WT-LRRK2 expressing cells

The LRRK2 kinase inhibitor EB-42168 was previously shown to inhibit G2019S LRRK2 100-fold more potently than wild-type LRRK2 [20]. MLi‐2, by contrast, was equipotent in recombinant cell lines expressing WT-LRRK2 or G2019S-LRRK2, and can therefore serve as a nonselective LRRK2 kinase inhibitor control [20, 24]. We assessed the potency of EB-42168 and MLi-2 to inhibit pSer935 and pSer1292 LRRK2 in WT or G2019S-LRRK2 expressing cells in compound titration experiments with concentrations ranging from 10 nM -1 μM. At the lowest concentration of MLi-2 tested (10 nM), pSer935 was significantly decreased by approximately 75% in both the WT-LRRK2 and G2019S-LRRK2 cells (Fig. 2A,B). At concentrations higher than 100 nM, MLi-2 showed nearly complete ablation of phosphorylation at Ser935, with <10% signal remaining (Fig. 2A,B). Overall, the concentration-response profile for MLi-2 at pSer935 was similar for the two genotypes (Fig 2 A,B). In comparison, EB-42168 inhibited pSer935 by over 50% at 100 nM with full inhibition at 1 μM (Fig 2C,D) but showed no inhibition of pSer935 in WT-LRRK2 cells up to 1 μM (Fig 2C,D).

**Fig 2.**
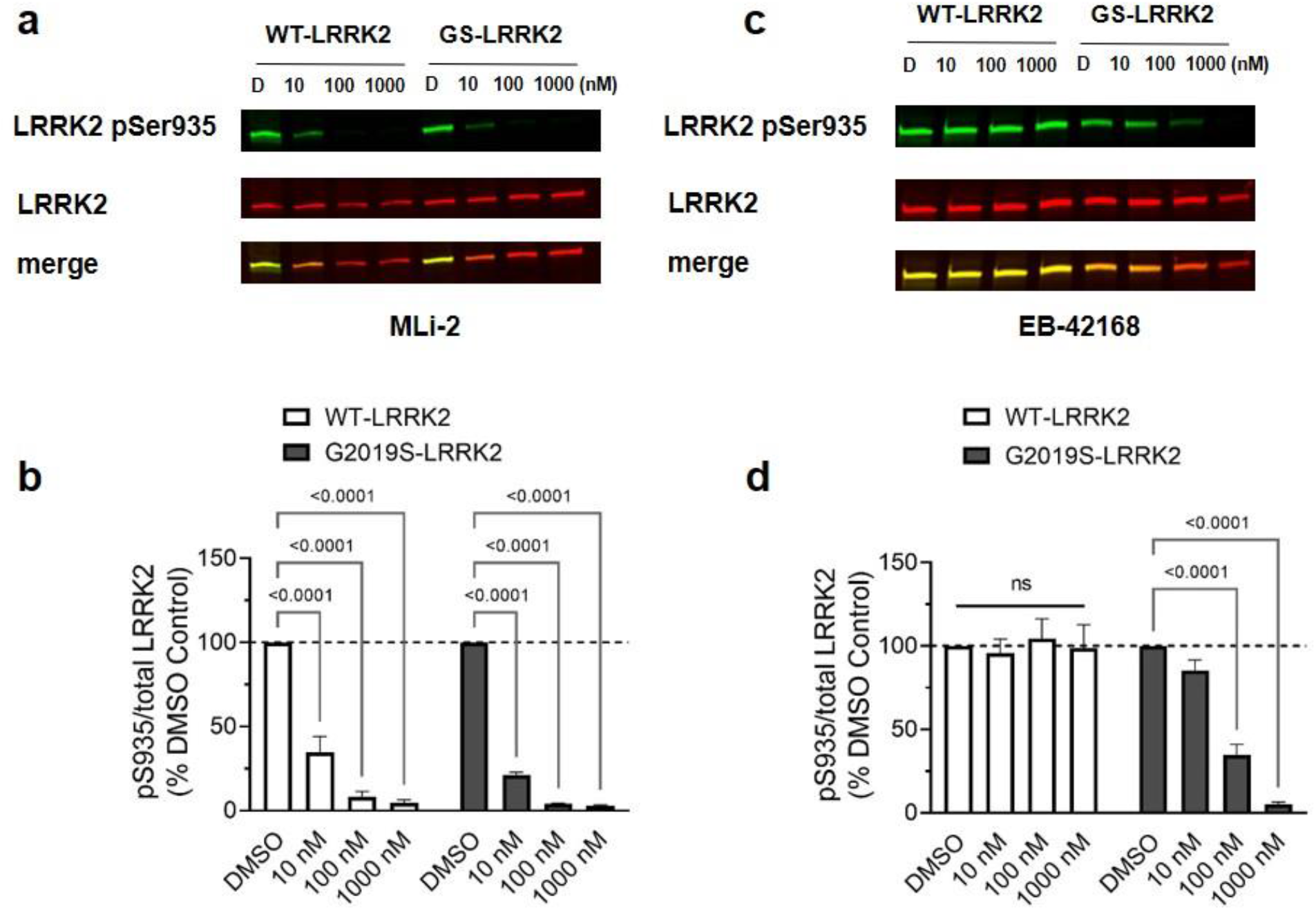
EB-42168 demonstrates full inhibition of LRRK2 pSer935 in cells overexpressing G2019S-LRRK2 at concentrations that show no inhibition in WT-LRRK2 overexpressing cells. **a** Representative western blot of WT-LRRK2 or G2019S-LRRK2 cells treated with DMSO, 10nM, 100nM or 1µM MLi-2 for 2h and assessed for LRRK2 pSer935 and full-length LRRK2. **b** Quantification of western blots demonstrate a 70%, 90% and 95% decrease respectively in LRRK2 pSer935 levels with increasing concentration of MLi-2 in WT-LRRK2 expressing cells. Quantification of western blots demonstrate an 80%, 96% and 97% decrease respectively in LRRK2 pSer935 levels with increasing concentration of MLi-2 in G2019S-LRRK2 expressing cells. **c** Representative western blot of WT-LRRK2 or G2019S-LRRK2 cells treated with DMSO, 10nM, 100nM or 1µM EB-42168 for 2h and assessed for LRRK2 pSer935 and full-length LRRK2. **d** Quantification of western blots demonstrate no change in LRRK2 pSer935 levels with exposure to EB-42168 in WT-LRRK2 expressing cells. Quantification of western blots demonstrate a 65% and 95% decrease respectively in LRRK2 pSer935 levels with 100nM and 1µM of EB-42168 in G2019S-LRRK2 expressing cells. Data are presented as mean ± SEM. (*P<0.01 determined by two-way ANOVA). G2019S-LRRK2 = GS-LRRK2, wild-type LRRK2 = WT-LRRK2, non-significant = ns

MLi-2 inhibited autophosphorylation at pSer1292 at a similar potency as pSer935 in G2019S-LRRK2 expressing cells. pSer1292 levels were significantly decreased following 10 nM treatment with MLi-2 (Fig 3A,B) and minimal pSer1292 levels remained at concentrations 100 nM and higher (Fig 3A,B). EB-42168 also showed similar potency on pSer1292 compared pSer935 at higher concentrations, with ∼70% reduction observed at 100 nM and full inhibition at 1 μM (Fig 3C,D). A modest decrease in pSer1292 was observed at 10nM. Due to the very low basal levels of pSer1292 in WT-LRRK2 cells, we were unable to accurately quantitate inhibition of the pSer1292 signal using the immunoblot method in the WT-LRRK2 cell line and therefore unable to assess genotype selectivity of the inhibitors on LRRK2 autophosphorylation.

**Fig 3.**
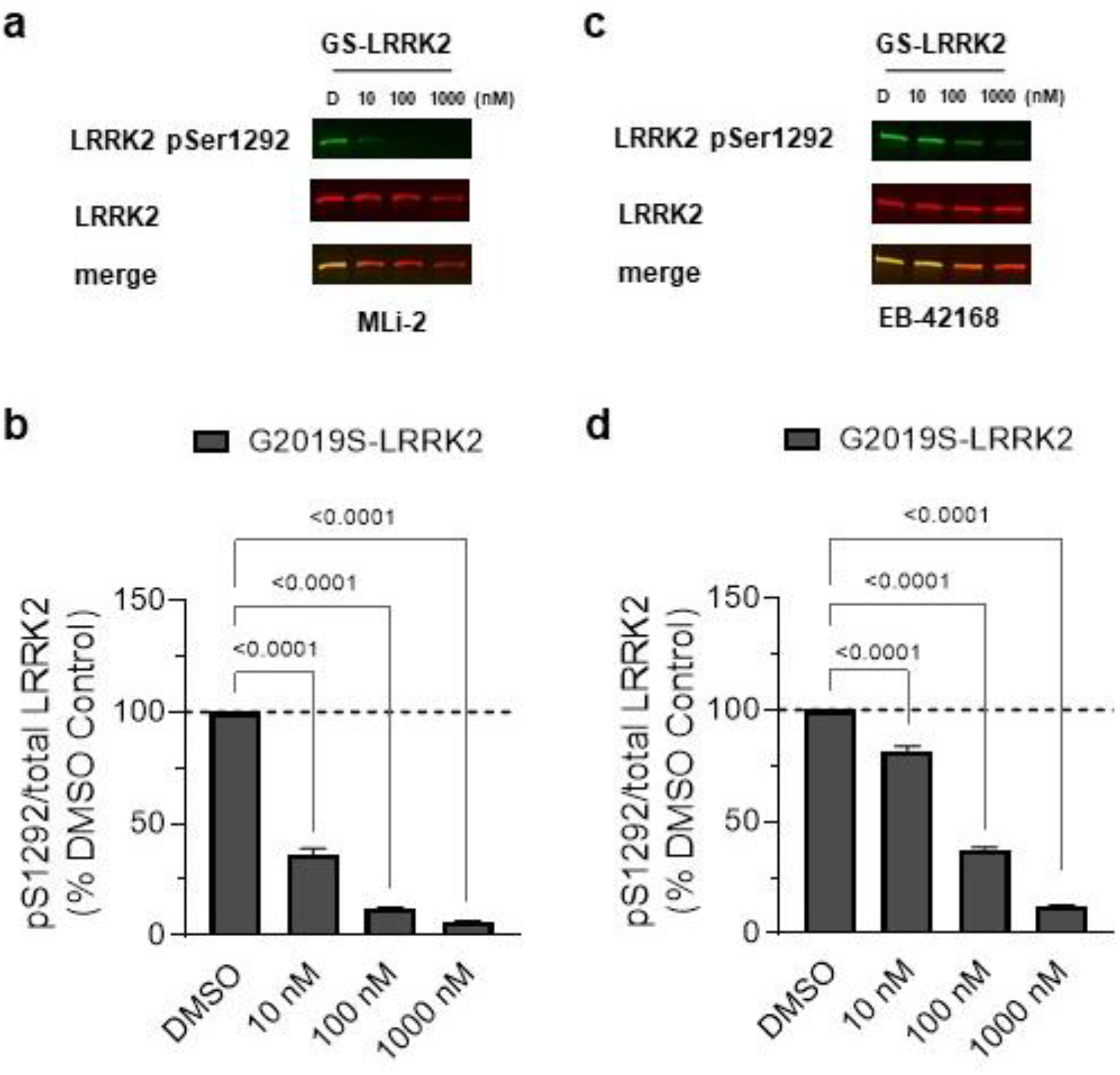
EB-42168 inhibits LRRK2 pSer1292 in G2019S-LRRK2 overexpressing cells at concentrations that also inhibited pSer935. **a** Representative western blot of G2019S-LRRK2 cells treated with DMSO, 10nM, 100nM or 1µM MLi-2 for 2h and assessed for LRRK2 pSer1292 and full-length LRRK2. **b** Quantification of western blots demonstrate a 65%, 90% and 95% decrease in LRRK2 pSer1292 levels with increasing concentration of MLi-2 in G2019S-LRRK2 expressing cells. **c** Representative western blot of G2019S-LRRK2 cells treated with DMSO, 10nM, 100nM or 1µM EB-42168 for 2h and assessed for LRRK2 pSer1292 and full-length LRRK2. **d** Quantification of western blots demonstrate a 30%, 65% and 85% decrease in LRRK2 pSer1292 levels with 10nM, 100nM and 1µM respectively of EB-42168 in G2019S-LRRK2 expressing cells. Data are presented as mean ± SEM. (*P<0.01 determined by one-way ANOVA). G2019S-LRRK2 = GS-LRRK2

### Acute exposure to G2019S-selective kinase inhibition mitigates G2019S LRRK2-induced mtDNA damage

The loss of constitutive phosphorylation of LRRK2 at Ser935 in the presence of a LRRK2 kinase inhibitor is rapid [[53, 54] (Fig. 2,3)]. We recently reported that acute exposure of G2019S LRRK2 patient-derived cells to LRRK2 kinase inhibitors restored mtDNA damage to control levels [43, 48]. To determine whether the time course of mtDNA damage reversal by LRRK2 kinase inhibitors also occurs quickly with a G2019S-selective inhibitor, WT-LRRK2 or G2019S-LRRK2 cells were exposed to MLi-2, EB-42168, or vehicle for 2h. Exposure of G2019S-LRRK2 cells to MLi-2 restored mtDNA damage to WT-LRRK2 control levels at concentrations ranging from 10 nM to 1 μM (Fig. 4A). Culturing in the presence of MLi-2 at higher concentrations induced a trend towards an increase in mtDNA damage in WT-LRRK2 cells (Fig 4A). MLi-2 had no effect on mtDNA copy number in WT-LRRK2 and G2019S-LRRK2 cells (Fig 4B). Interestingly, all concentrations of EB-42168 tested alleviated G2019S-LRRK2 induced mtDNA damage, normalizing damage to WT-LRRK2 control levels (or lower) within 2 hours of compound addition (Fig 4C). No effect of EB-42168 was observed in WT-LRRK2 cells at the range of concentrations tested (Fig 4C). Mitochondrial DNA copy number was not different in G2019S-LRRK2 or WT-LRRK2 cells following an acute exposure of EB-42168 (Fig. 4D).

**Fig 4.**
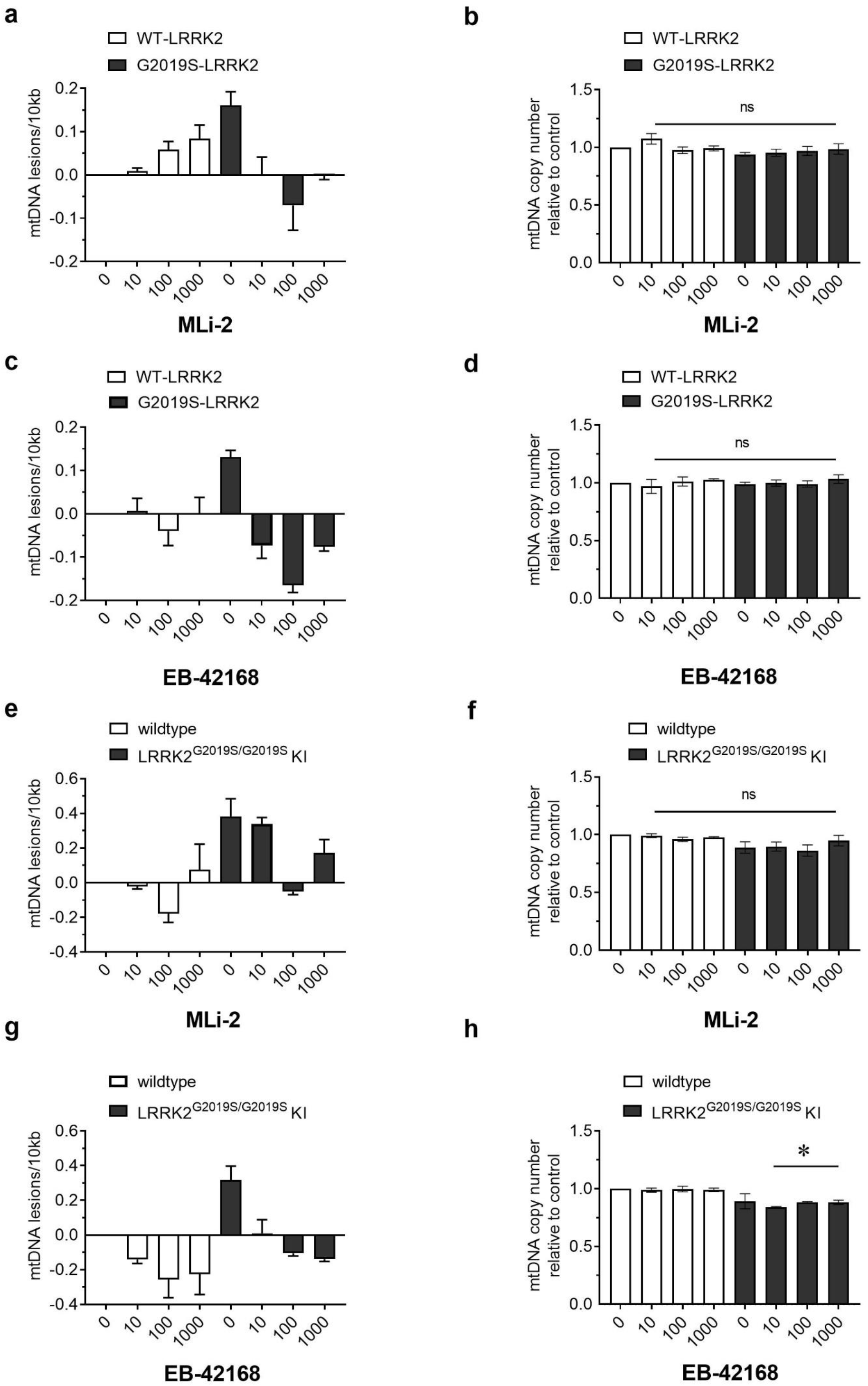
Mitochondrial DNA damage in G2019S-LRRK2 overexpression and LRRK2^G2019S/G2019S^ KI cells was abrogated to normal levels with LRRK2 kinase inhibition. **a** WT-LRRK2 and G2019S-LRRK2 cells were treated with DMSO, 10nM, 100nM or 1µM MLi-2 or **c** EB-42168 for 2h and analyzed for mtDNA damage. **b** and **d** mtDNA copy number was unaltered by these treatments. **e** LRRK2^G2019S/G2019S^ KI cells and wild-type cells were treated with DMSO, 10nM, 100nM or 1µM MLi-2 or **g** EB-42168 for 2h and analyzed for mtDNA damage. **f** MLi-2 exposure did not change mtDNA copy number in either cell line. **h** In contrast, while EB-42168 had no effect on mtDNA copy number in wild-type cells, mtDNA copy number was modestly decreased in LRRK2^G2019S/G2019S^ KI cells. Data are presented as mean ± SEM. (*P<0.01 determined by one-way ANOVA). G2019S-LRRK2 = GS-LRRK2, wild-type LRRK2 = WT-LRRK2, non-significant = ns

To investigate the effect of MLi-2 and EB-42168 in cells with endogenous levels of LRRK2, we utilized homozygous G2019S knock-in (LRRK2^G2019S/G2019S^ KI) HEK293 cells generated by CRISPR/Cas9 gene editing [32]. Mitochondrial DNA damage was statistically significantly increased in LRRK2^G2019S/G2019S^ KI cells compared to wild-type cells (Fig. 4E), without a change in mtDNA copy number (Fig. 4F). In LRRK2^G2019S/G2019S^ KI cells, a 100 nM concentration of MLi-2 reversed mtDNA damage levels to wild-type levels (Fig. 4G). No effect was observed on mtDNA copy number (Fig. 4H). In contrast, mtDNA damage in LRRK2^G2019S/G2019S^ KI cells were reversed to wildtype levels at all EB-42168 concentrations tested (Fig. 4I). Interestingly, mtDNA copy number was reduced in EB-42168 treated LRRK2^G2019S/G2019S^ KI cells (Fig. 4J).

### EB-42168 demonstrates selective inhibition in heterozygous G2019S LRRK2 PD patient-derived cells compared to healthy controls

To investigate the selective effect of EB-42168 relative to MLi-2 on LRRK2 phosphorylation and mtDNA damage biomarkers in PD patient cells, LRRK2 G2019S and healthy control derived lymphoblastoid cells (LCLs) were evaluated. Of note, only LRRK2 pSer935 levels were measured in response to LRRK2 kinase inhibition, as the pSer1292 signal in LCLs was not quantifiable by immunoblotting methods [11, 48]. MLi-2 at 10 and 100 nM reduced pSer935 almost completely in both healthy controls and G2019S LRRK2 PD patient-derived LCLs, similar to previously published results (Fig 5 A,B [48]). EB-42168 exposure did not impact pSer935 levels in healthy controls (Fig 5 C,D). In contrast, in heterozygous G2019S LRRK2 PD patient-derived LCLs, treatment with EB-42168 partially reduced LRRK2 pSer935 phosphorylation, ∼30 and 40% with 10 and 100 nM, respectively (Fig 5 C,D), presumably due to a lack of sensitivity on WT LRRK2 which accounts for ∼50% of the total pSer935 signal in heterozygous cells.

**Fig 5.**
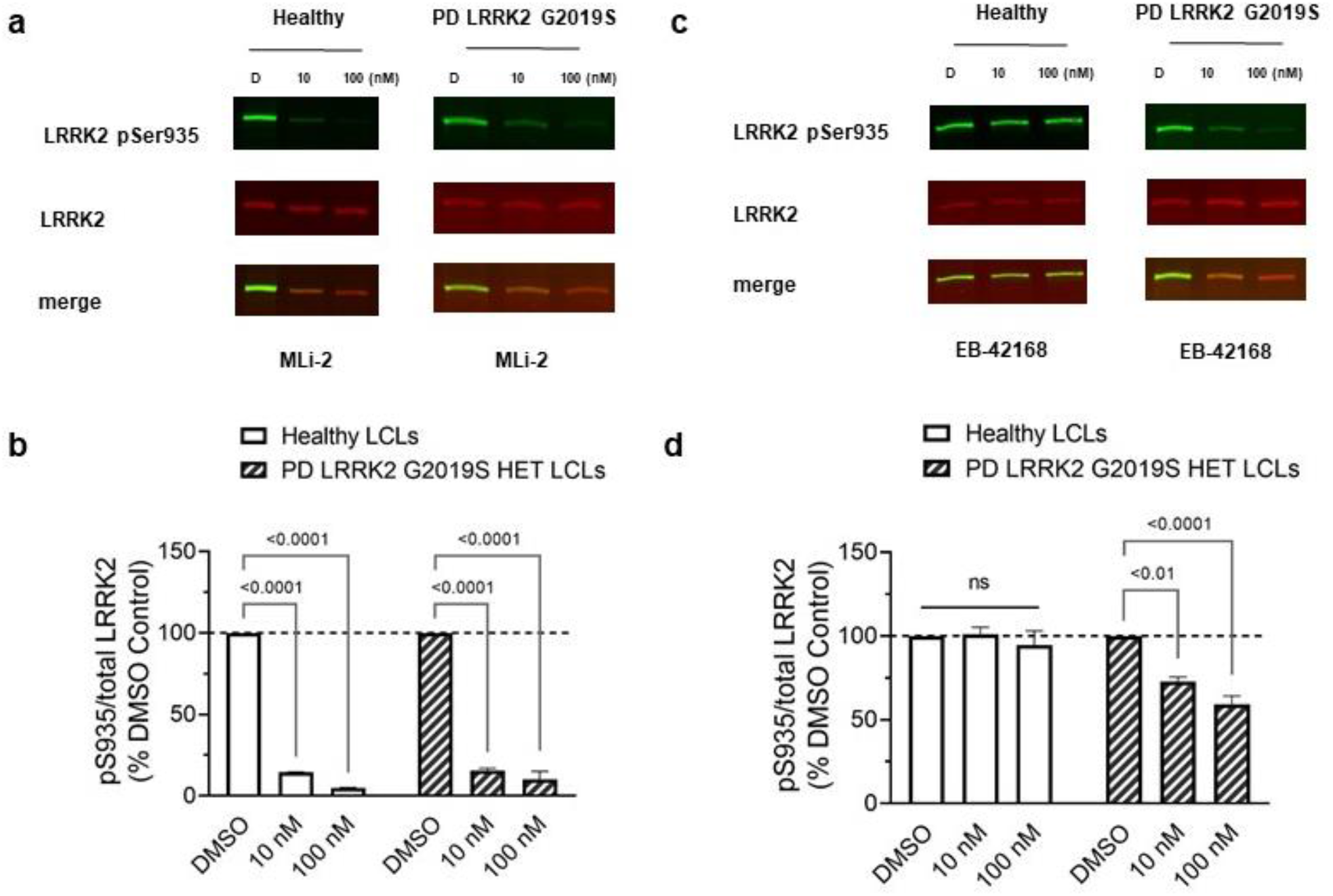
EB-42168 demonstrates selective inhibition of pSer935 LRRK2 in heterozygous G2019S LRRK2 PD-patient derived LCLs relative to healthy controls. **a** Representative western blot of PD patient-derived LRRK2 G2019S carriers and age-matched healthy controls LCLs treated with DMSO, 10nM, or 100nM MLi-2 for 2h and assessed for LRRK2 pSer935 and full-length LRRK2. **b** Quantification of western blots demonstrate a 85% and 95% decrease in LRRK2 pSer935 levels in LRRK2 G2019S PD patient-derived and age-matched healthy controls. **c** Representative western blot of PD patient-derived LRRK2 G2019S carriers and age-matched healthy controls LCLs treated with DMSO, 10nM, or 100nM EB-42168 for 2h and assessed for LRRK2 pSer935 and full-length LRRK2. **d** Quantification of western blots demonstrate no change in LRRK2 pSer935 levels with exposure to EB-42168 in healthy controls. Quantification of western blots demonstrate a 25% and 40% decrease in LRRK2 pSer935 levels with 10nM and 100nM EB-42168 in PD patient-derived LRRK2 G2019S carriers. Data are presented as mean ± SEM. (*P<0.01 determined by two-way ANOVA).

Mitochondrial DNA damage levels were measured in parallel. Acute treatment with MLi-2 similarly reversed mtDNA damage in G2019S LRRK2 patient-derived LCLs at all concentrations tested, but had no effect in healthy controls (Fig 6A). No differences in mtDNA copy were detected with MLi-2 treatment, regardless of PD status or genotype (Fig. 6B). Importantly, similar results were found with EB-42168: mtDNA damage was restored to control levels at concentrations of 10 and 100 nM in heterozygous G2019S LRRK2 LCLs, without an effect in healthy controls (Fig 6C). No differences in mtDNA copy were detected with EB-42168 treatment in either healthy controls or PD patient-derived G2019S LRRK2 LCLs (Fig 6D).

**Fig 6.**
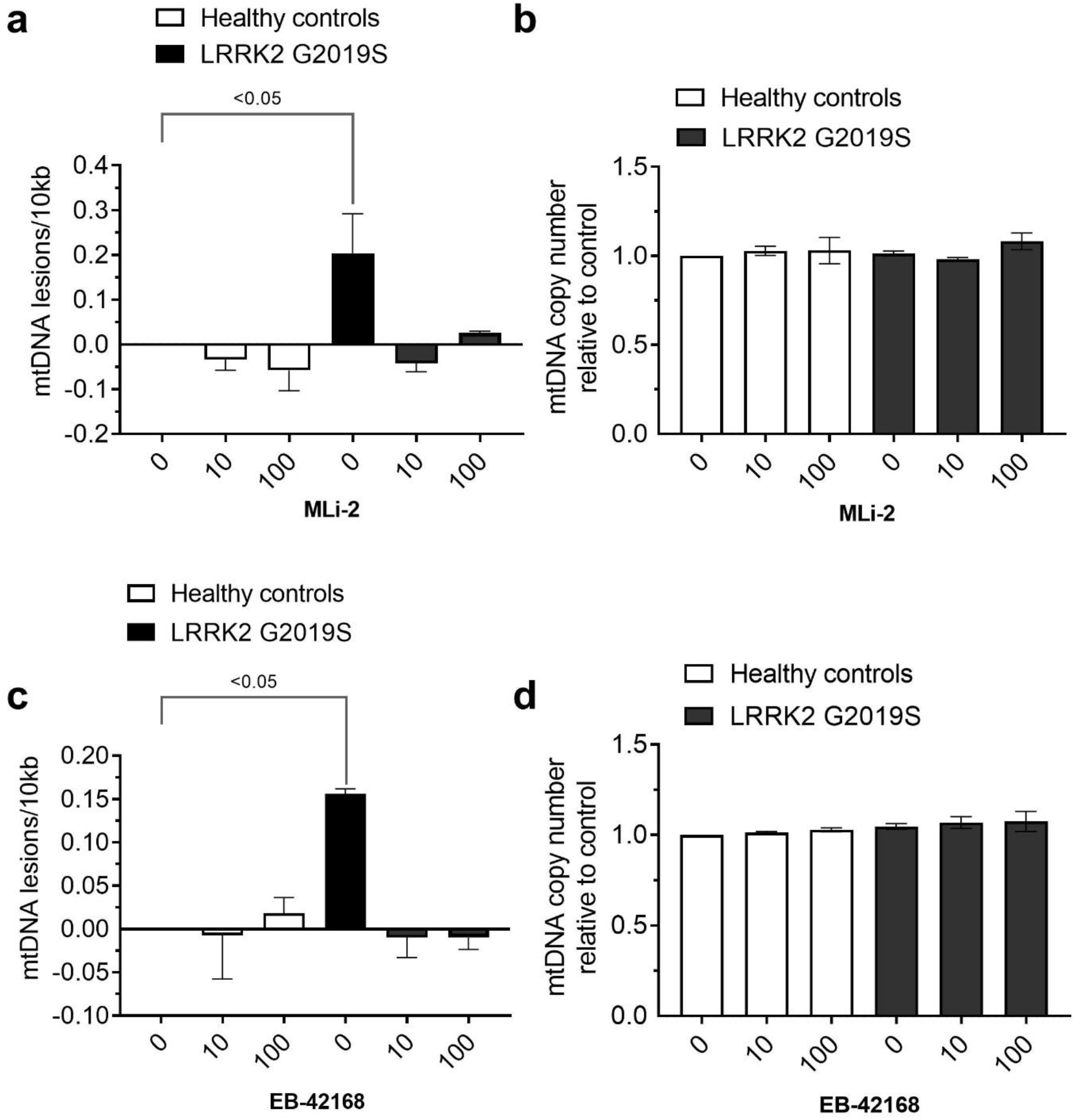
LRRK2 kinase inhibitor exposure restored LRRK2 G2019S induced mtDNA damage to basal healthy control levels. **a** PD patient-derived LRRK2 G2019S carriers and age-matched healthy controls LCLs treated with DMSO, 10nM, or 100nM MLi-2 or **c** EB-42168 for 2h and analyzed for mtDNA damage. **b** and **d** mtDNA copy number was unaltered by these treatments. Data are presented as mean ± SEM. (*P<0.01 determined by one-way ANOVA).

## DISCUSSION

Significant progress has been made elucidating cellular pathways that are altered by kinase-activating LRRK2 variants associated with familial and idiopathic PD. Previously, we demonstrated that mtDNA damage is increased specifically in dopaminergic neurons in human PD patient-derived post-mortem brains and *in vivo* PD models [42]. Mitochondrial DNA damage is elevated in PD patient-derived LRRK2 mutant neurons and immune cells and is dependent on LRRK2 kinase activity [42-44, 48]. We further demonstrated that mtDNA damage in human heterozygous G2019S LRRK2 PD patient-derived cells was restored to healthy control levels following exposure to LRRK2 kinase inhibitors in a dose-dependent fashion [48]. The objective of the current study was to investigate whether G2019S LRRK2 dependent mtDNA damage could also be abrogated by selectively inhibiting only pathogenic LRRK2 kinase activity derived from the G2019S LRRK2 mutant allele. Our data demonstrates that mtDNA damage is induced by the PD-associated G2019S mutation in LRRK2, either in an overexpressed context or KI model, thus indicating for the first time that the G2019S mutant expressed at endogenous levels is sufficient to drive mtDNA damage. We further illustrate that the mtDNA damage phenotype can be restored by pharmacological treatments with both selective and non-selective LRRK2 inhibitors in LRRK2 cellular models and G2019S LRRK2 PD patient-derived cells. Together, these data indicate that mtDNA damage can serve as a functional pharmacodynamic marker for G2019S-selective and non-selective LRRK2 kinase inhibition and provide critical insight into LRRK2 based therapeutics.

*LRRK2* pathogenic mutations, including the G2019S variant, lead to hyperactive kinase activity [6-8]. As such, LRRK2-associated PD medicinal chemistry programs have focused on the development of LRRK2 kinase inhibitors. While these inhibitors have high specificity for the kinome, most do not discriminate between WT-LRRK2 and pathogenic G2019S-LRRK2. The strategy of pursuing specific G2019S-LRRK2 kinase inhibitors may be critical in mitigating potential safety liabilities associated with the impact of non-selective LRRK2 kinase inhibitors on the wild-type allele. In this pursuit, the first identification of small molecules that could precisely target a mutation in LRRK2 was discovered in a high-throughput screen, of which the scaffold was further optimized for a series of novel, potent compounds that preferentially inhibited G2019S-LRRK2 [20]. Utilizing one of these compounds, EB-42168, we found with increasing concentrations that LRRK2 dephosphorylation was only observed in G2019S-LRRK2 expressing cells with no detectable effect on WT-LRRK2. These results are consistent with previous findings that EB-42168 inhibited G2019S-LRRK2 100-fold more potently than WT-LRRK2 [20]. In human PD-patient derived cells carrying a heterozygous G2019S LRRK2 mutation, we found that LRRK2 pSer935 was decreased about 40%, without a change in healthy controls. Importantly, the effect of EB-42168 was similar in an artificial overexpression system and in human derived cells with endogenous levels of human LRRK2. Similar levels of inhibition of pSer935 LRRK2 with EB-42168 in heterozygous G2019S LRRK2 carriers derived from human peripheral blood mononuclear cells has been reported [50]. On the other hand, a non-selective inhibitor, MLi-2 at 10nM and above, demonstrated almost complete pSer935 LRRK2 inhibition, in both G2019S and WT-LRRK2 overexpression systems and in human derived healthy controls and G2019S LRRK2 PD-patient cells; not surprisingly based on previous findings [43, 48, 50]. Although EB-42168 is a poorly brain penetrant molecule [20], brain penetrant G2019S-selective LRRK2 inhibitors with promising pharmacokinetic properties have been identified [51, 55]. Testing brain penetrant G2019S-LRRK2 selective compounds *in vivo* on LRRK2 phosphorylation and mtDNA damage biomarkers will be explored in future studies.

While the rationale for targeting pathogenic LRRK2 is compelling, the precise amount of kinase inhibition that is needed to be potentially efficacious remains to be elucidated. This is partially due to the inherent challenges in using surrogate markers of activity. Current potential brain and peripheral biomarkers of LRRK2 kinase activity are either indirect, lack sensitivity or do not correlate with intrinsic cellular kinase activity of LRRK2, posing challenges to the development of LRRK2-targeted therapeutics [35, 56-60]. It is also unclear that the current biomarkers can capture the dynamic nature of LRRK2 kinase activity and whether brain and peripheral changes are equivalent comparisons. We previously found with non-selective LRRK2 kinase inhibitors that ∼ 25% inhibition of LRRK2 pSer935 was sufficient to reverse mtDNA damage levels back to healthy control levels [48]. However, we were not able to reliably measure pSer1292 LRRK2 under the same conditions, a barrier in most endogenous contexts [31]. An advantage to the current study is that we utilized stably overexpressing LRRK2 cell lines to compare how pSer1292 and pSer935 LRRK2 dephosphorylation changes with non-selective and selective LRRK2 kinase inhibitors. At the MLi-2 concentrations tested, LRRK2 pSer935 and pSer1292 had similar trends and equal levels of dephosphorylation. However, to our surprise, at 10nM of EB-42618, no significant dephosphorylation of LRRK2 pSer935 was detected, yet LRRK2 pSer1292 was dephosphorylated, indicating that LRRK2 kinase activity was inhibited. Even without target engagement of pSer935 LRRK2 at 10nM with EB-42618 by immunoblotting, mtDNA damage was abrogated, highlighting that mitochondrial genome integrity is extremely sensitive and may accurately reflect intrinsic LRRK2 kinase activity [12, 39, 40]. The majority of preclinical studies have evaluated neuroprotection or related PD-pathology with >90% pSer935 inhibition [61]. Yet, only partial inhibition of LRRK2 kinase activity could be sustained without lung pathology in monkeys [22]. Therefore, future preclinical and clinical studies may consider evaluating these multiple biomarkers, and levels of mtDNA damage in human blood hold promise for guiding this balancing act of efficacy and minimizing safety risk [62].

Under conditions in which G2019S-LRRK2 and WT-LRRK2 are expressed equally, we found that mtDNA damage levels are significantly enhanced with the PD mutation. These results are consistent with our previous findings that expression of WT-LRRK2 had no effect on mtDNA damage in neurons [43]. These data suggest that accumulating mtDNA damage depends on LRRK2 kinase activity and not LRRK2 levels. Alternative approaches to therapeutically targeting LRRK2 have emerged [13]. Antisense oligonucleotides (ASOs) in a preclinical proof of concept study showed that LRRK2 mRNA was reduced in the brain while sparing peripheral tissues [63]. Early phase human clinical studies are underway using an ASO targeting LRRK2 (ClinicalTrials identifier NCT03976349). The on-going and future ASO and LRRK2 kinase inhibitor targeted approaches will examine if reducing LRRK2 levels and/or kinase activity ameliorates the toxic gain-of-function effects of LRRK2 in PD patients and results in clinically relevant efficacy.

A precision medicine approach is common in many fields, but has only been recently applied to neurodegenerative diseases and could in part explain the failures of this “one size fits all” approach to PD trials [64]. A precision medicine approach that accounts for an individual’s genotype and/or phenotype may pave the way for successful disease-modifying treatments. Although, this approach is not without its challenges and does not guarantee a positive outcome (ClinicalTrials.gov Identifier: NCT02906020). Biomarkers will be critical to guide patient stratification, response to treatment and disease progression. Advancing a precision medicine based therapeutic approach for LRRK2-associated PD will require integrated measures with a range of clinical and molecular parameters [65].

## Materials and Methods

### LRRK2 cell lines and PD patient *LRRK2* mutation carriers and healthy control lymphoblastoid cells

HEK293 cells stably transfected with human LRRK2 or the G2019S variant of human LRRK2 (WT-LRRK2 and G2019S-LRRK2, [20] were maintained in an incubator at 37°C with 5% CO_2_ and grown in Eagle’s Minimum Essential Medium (ATCC: The Global Biosource, 30-2003), 10% Gibco Fetal Bovine Serum, Qualified (Fisher Scientific, 10-437-028), and 0.5% Penicillin Streptomycin (Corning, 30-002-Cl). Geneticin (Thermo Fisher Scientific, 10131035) was added to media with cells at a concentration of 400 μg/mL. Cells were plated at a density of 0.5 × 10^6^ in a 6-well dish and treated at ∼ 40-50 % confluency. LRRK2^G2019S/G2019S^ KI HEK293 cells [32] were maintained in DMEM/F12, GlutaMAX™ supplement (Gibco, 10565-018), Seradigm Premium Grade 10% FBS (VWR, 97068-085) and 0.5% Penicillin Streptomycin (Corning, 30-002-Cl). Cells were plated at a density of 0.3 × 10^6^ cells in a 6-well dish and treated at ∼ 40-50 % confluency. At the time of harvest, cells were collected and either protein collected or DNA extracted as described below.

PD patient G2019S LRRK2 (*n =* 2) and healthy subject control (*n =* 2)-derived LCLs were obtained from the NINDS Coriell biorepository (ID numbers are ND00011, ND00264, ND01618, ND02559). There was not a statistical difference in the ages between the PD patient and healthy control subjects (*P* = 0.92). LCLs were cultured at 37°C, 5% CO_2,_ in RPMI-1640 (Sigma-Aldrich, R8758), 15% heat-inactivated fetal bovine serum (VWR Seradigm, 97068-091) and 0.5% Penicillin/Streptomycin (Corning, 30-002-CI). Cells were passaged every 3-4 days, and passage number did not exceed 20.

### LRRK2 kinase inhibitors

Multiple LRRK2 kinase inhibitors were utilized for *in vitro* experiments including MLi-2 and EB‐42168 [20, 24]. The tool compound EB-42168 was discovered as part of a small molecule discovery program aimed at identifying LRRK2 kinase inhibitors that are selective for the pathogenic G2019S LRRK2 variant [20]. MLi‐2 (synthesized at WuXi in Tianjing, China), was included as a reference inhibitor and was shown previously to exhibit similar binding affinity for both WT and G2019S LRRK2 [24]. MLi‐2 inhibits WT LRRK2 approximately 1000‐fold more than EB‐42168 [20]. For all compound treatments, cells were treated for 2 hours with varying concentrations of EB-42168 or MLi-2. Compounds were dissolved in DMSO at an initial concentration of 20 mM and serially diluted in DMSO and added to media to achieve a final concentration in the range of 10 nM -1 μM and the same final DMSO concentration for each solution.

### Western blot analysis and antibodies

For LRRK2-WT or G2019S-LRRK2, ∼1.5 million cells were pelleted and resuspended in 75μl of lysis buffer [(RIPA Buffer (Sigma-Aldrich, R0278-50ML), protease inhibitor cocktail (Sigma-Aldrich, P8340), and Halt phosphatase inhibitor cocktail (Thermo Fisher, 78420)]. For LCLs, five million cells were pelleted and resuspended in 100 µl of lysis buffer. For both types of cells lysates were centrifuged at 10,000 × *g* after a 10 minute incubation on ice, and the supernatant was collected. Protein was quantified using the DC protein assay (Bio-Rad, 5000112). 50 μg of protein was loaded for HEK293 LRRK2-WT or G2019S-LRRK2 cells. Due to the differing endogenous LRRK2 levels, 40 µg of ND02559, 60 µg of ND00264, or 100 µg of ND00011 and ND01618 of protein sample was loaded. Protein samples were incubated at 100°C for 5 min with NuPAGE Sample loading dye (Thermo Fisher, NP0007) and dithiothreitol as reducing agent. After 4-20% SDS-PAGE, the blots were blocked in 5% w/v nonfat dry milk in 1X PBST (0.05% Tween 20). For our investigations, the following primary antibodies were used: mouse anti-LRRK2 N241a (Antibodies INC, 75-253, 1:5000 for HEK293 cells and 1:2000 for LCLs), rabbit anti-LRRK2 pS935 (Abcam, ab133450, 1:10,000 for HEK293 cells and 1:2000 for LCLs), mouse anti-β-actin (Novus, 8H10D10, 1:10,000). rabbit anti-LRRK2 pS1292 (Abcam, ab203181, 1:500). The blots were then probed with fluorescent-labeled secondary antibodies, IRDye donkey anti-mouse and anti-rabbit at 1:10,000 (LI-COR, 926-32212, 926-68072), and scanned using an Odyssey Imaging scanner (LI-COR). Fluorescence intensities were quantified using ImageStudio Lite software (LI-COR), and the signal from the protein of interest was normalized to the fluorescence intensity of either LRRK2 or β-actin. Values were averaged from at least three technical replicates within a cell line, or two biological replicates (two healthy control lines and two PD LCLs).

### DNA isolation, quantitation and mitochondrial DNA damage measurement

Cells were collected and the nuclei and mitochondria were isolated as previously described [48]. DNA was extracted from LRRK2-WT and G2019S-LRRK2 cells, and wildtype and LRRK2^G2019S/G2019S^ KI cells using QuickGene DNA Whole Blood Kit L (Autogen, fk-dbl). PD patient LRRK2 G2019S and healthy subject control -derived LCL DNA was extracted using the QuickGene DNA Tissue Kit L (Autogen, fk-dtl), utilizing a semi-automated system (Autogen, QuickGene-610L). For the Autogen system, the pellet was resuspended in 2 mL of 1X PBS, and the standard manufacturer’s protocol was conducted as if the cell suspension was whole blood. DNA was eluted with EDTA-free buffer, and quality was assessed using a Spectradrop microvolume microplate (Molecular Devices). Double-stranded DNA was quantified using Quant-iT Picogreen dsDNA assay (Thermo Fisher) as previously described [48].

### PCR-based mitochondrial DNA damage assay

DNA damage in the mitochondrial genome was measured utilizing Mito DNA_DX_ [62], currently the most robust way of measuring damage in mtDNA [66]. The assay to calculate mitochondrial DNA lesion frequency was performed as previously described [62]. Briefly, 15 ng of DNA was used to amplify long or short amplicons of the mitochondrial genome (as determined by primer sets). The amount of amplification is directly proportional to the number of undamaged DNA templates. Average lesion frequency is calculated as described [62]. Results are then normalized and presented as lesions per 10 kb, with the values in experimental samples relative to control samples. PCR reactions included KAPA Long Range HotStart DNA Polymerase (KAPA Biosystems) [48]. Each biological DNA sample was analyzed in technical triplicate.

### Statistical analyses

Data were analyzed in Prism software (GraphPad). Data were analyzed by either unpaired, two-tailed Student t-test or ANOVA with a Tukey or Bonferroni’s post-hoc analysis. *P*-values < 0.05 were considered significant. For all graphs, the bars represent mean ± standard error of the mean (SEM).

## Acknowledgements

We would like to acknowledge Gretchen Argast for her input in the original study design. We are grateful to Tim Greenamyre for sharing the LRRK2^G2019S/G2019S^ KI cells. We also acknowledge funding support from ESCAPE Bio (L.H.S), and NINDS R01NS119528 (L.H.S.)

## Authors’ Roles

H.C. and L.H.S designed the project. N.P, C.P.G., R.Q., H.C., and L.H.S. performed experiments and data analysis. N.P., H.C., and L.H.S. wrote the manuscript. All authors read and approved the final manuscript.

